# A computational model for quantifying instability of tandem repeats across the genome

**DOI:** 10.64898/2026.04.08.717199

**Authors:** Egor Dolzhenko, Adam English, Tom Mokveld, Guilherme de Sena Brandine, Zev Kronenberg, Galen Wright, Britt Drögemöller, William J. Rowell, Aaron M. Wenger, Xiao Chen, Mark F. Bennett, Ben Weisburd, Graham S. Erwin, Peng Jin, David Nelson, Harriet Dashnow, Fritz J. Sedlazeck, Michael A. Eberle

## Abstract

Tandem repeats (TRs) exhibit high levels of somatic mosaicism, which is increasingly recognized as an important modifier of repeat expansion disorders. Long-read sequencing can capture full-length repeat alleles, yet robust frameworks for quantifying instability across TRs genome-wide are still needed. Here, we introduce a general-purpose model for quantifying TR instability in a given long-read sequencing dataset, without explicitly distinguishing biological mosaicism from technical noise, and which is broadly applicable to both simple and structurally complex loci. This model accurately characterizes allelic instability at each TR locus by representing the distribution of read-to-consensus deviations for each allele. Using HiFi sequencing data from 256 HPRC cell line samples, we fitted models for 617,007 TR loci, including known pathogenic repeats. We observe that instability levels are generally low, but vary substantially across individual TRs, and are driven more strongly by repeat composition than overall repeat length. Furthermore, we applied our method to targeted PureTarget long-read data from samples with known repeat expansions and identified significant mosaicism in the majority of expanded alleles. Our model offers a practical way to quantify instability of tandem repeats across the genome and to detect unusually unstable repeat alleles.

## Introduction

Tandem repeats (TRs) are among the most mutable genomic regions, and repeat mosaicism has been linked to the age of onset in repeat expansion disorders. In Huntington disease, for example, somatic instability is associated with earlier onset and faster disease progression, supporting a direct role for mosaicism in disease pathogenesis and generating strong interest in quantifying repeat instability genome-wide (1,2). This has positioned somatic instability as a candidate biomarker for clinical trials in repeat expansion disorders, while also motivating the development of therapeutic strategies that target these processes (3). Advances in the accuracy of long-read sequencing data have made it possible to fully resolve TR alleles that are too complex to reconstruct from short-read data, enabling direct measurement of heterogeneity in repeat length and sequence composition (4). Yet computational approaches for modeling this variability genome-wide remain limited, making it difficult to characterize the baseline instability of a repeat region or to identify highly unstable alleles.

We present a general-purpose method for quantifying TR instability in long-read sequencing data across a cohort of samples. The method measures read-to-consensus divergence for each allele and uses this information to model the baseline instability expected at each repeat locus in a target population. This provides a practical way to compare instability across repeats and to identify alleles whose instability is unusually high relative to their repeat-specific baseline. In practice, such alleles may be prioritized for follow-up as candidate pathogenic variants, particularly at loci already associated with repeat expansion disorders.

Because precise orthogonal assessments of mosaicism are generally unavailable, it is often difficult to distinguish read-level heterogeneity arising from true biological mosaicism from that introduced by technical noise. We therefore do not attempt to separate these sources of variation here. Instead, we focus on modeling the overall instability observed in long-read sequencing data, allowing us to define a repeat-specific baseline against which unusually unstable alleles can be detected.

Applying our model to 617,007 TR regions across 256 HiFi whole-genome sequencing cell line samples from the Human Pangenome Reference Consortium (5–7), we found that per-read instability rates are generally low but vary substantially across loci. Instability rates across repeats had a stronger association with repeat purity than with repeat length, confirming repeat sequence composition as a major driver of TR instability. Analysis of targeted long-read sequencing data from cell line samples with known repeat expansions revealed that the expanded alleles were significantly more unstable than the repeat-specific baseline. Taken together, our results establish a general-purpose framework for quantifying instability of TRs and for detecting and monitoring abnormally unstable alleles that are often linked to pathogenic expansions.

## Results

We developed a general-purpose method to measure repeat instability (**Figure 1**) that is applicable to both simple and structurally complex TRs across the genome, including loci whose motif structure within the general population may not be fully resolved (e.g., *RFC1*). The method fits long-read sequencing data to a probabilistic model that describes instability of a given repeat across a cohort of samples and identifies alleles whose instability significantly exceeds the baseline.

**Figure 1.**
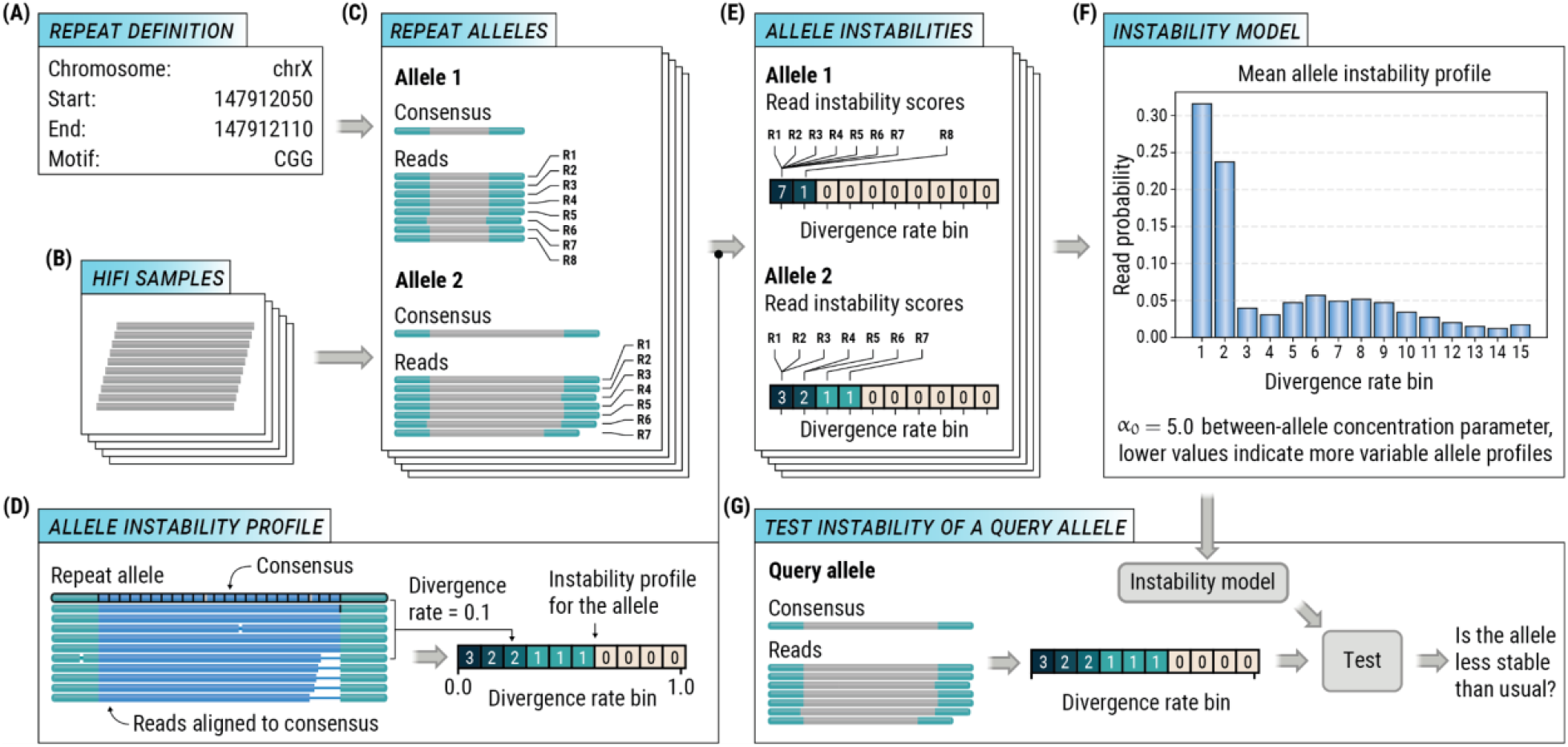
Overview of tandem repeat instability profiling. (A) Definition of a tandem repeat locus. (B) A cohort of HiFi samples was used to estimate baseline repeat instability. (C) For each locus, TRGT identifies the full-length consensus sequence of each repeat allele together with the reads supporting that allele. (D) For each supporting read, a divergence rate is computed as the length-normalized edit distance to the consensus allele sequence, then these read-level rates are binned to generate an allele instability profile. (E) Instability profiles are collected for all alleles of the repeat across the cohort. (F) A Dirichlet-multinomial model is fitted to these allele instability profiles to capture the baseline instability distribution of the repeat. (G) Instability of a query allele is assessed by comparing its profile to the fitted repeat-specific model.

To model the instability of TRs (**Figure 1A**) in a cohort of samples (**Figure 1B**), we first used our previously developed TR genotyping tool TRGT (4) to determine the full-length consensus sequence and supporting reads for every repeat allele. We then calculated a divergence rate for each read as the length-normalized edit distance between that read and its consensus allele sequence (**Figure 1C**). For each allele, we binned the divergence rates into 15 quantile-defined bins to generate an instability profile of read-counts per bin (**Figures 1D,E; Methods**). Finally, we estimated a Dirichlet-multinomial (DM) distribution for each repeat locus from the instability profiles of all observed alleles across the cohort, producing a locus-level model of expected instability (**Figure 1F**). In this model, the multinomial component describes variation in read depths across alleles, while the Dirichlet component captures overdispersion in instability profiles across a given TR’s alleles. To identify unusually unstable alleles, we compared each allele against its own TR model using a likelihood test statistic and assessed statistical significance using parametric bootstrap (**Figure 1G**; **Methods**).

### Properties of repeat instability rates

Read divergence rates form the basis of our model. We characterized these rates as a repeat instability rate by calculating the average divergence rate across all reads assigned to all alleles of a repeat across 617,007 TR regions from 256 HPRC whole-genome HiFi samples (**Methods**). Overall, read divergence rates were low with a mean repeat instability rate of 0.0051 and a median of 0.0039, indicating high concordance between consensus allele sequences and their reads. Because perfect repeats are known to be more unstable than interrupted repeats (8,9), we examined this relationship in our data. We defined repeat purity for each allele as the fraction of bases contained within its longest perfect repeat tract (10), and assessed the association of repeat instability rate with mean purity and mean allele length (**Methods**). Consistent with this expectation, repeat purity was strongly correlated with repeat instability rate (**Figure 2A**). By contrast, instability rate was only weakly correlated with overall allele length (Spearman’s rho = 0.09; **Figure 2B**), with the number of deviations between reads and the consensus allele sequence increasing approximately linearly with repeat length. This weak dependence on length is also consistent with the observation that perfect repeats in the human genome are typically short **(Figure 2C**), whereas longer alleles tend to contain interruptions (**Figure 2D**).

**Figure 2.**
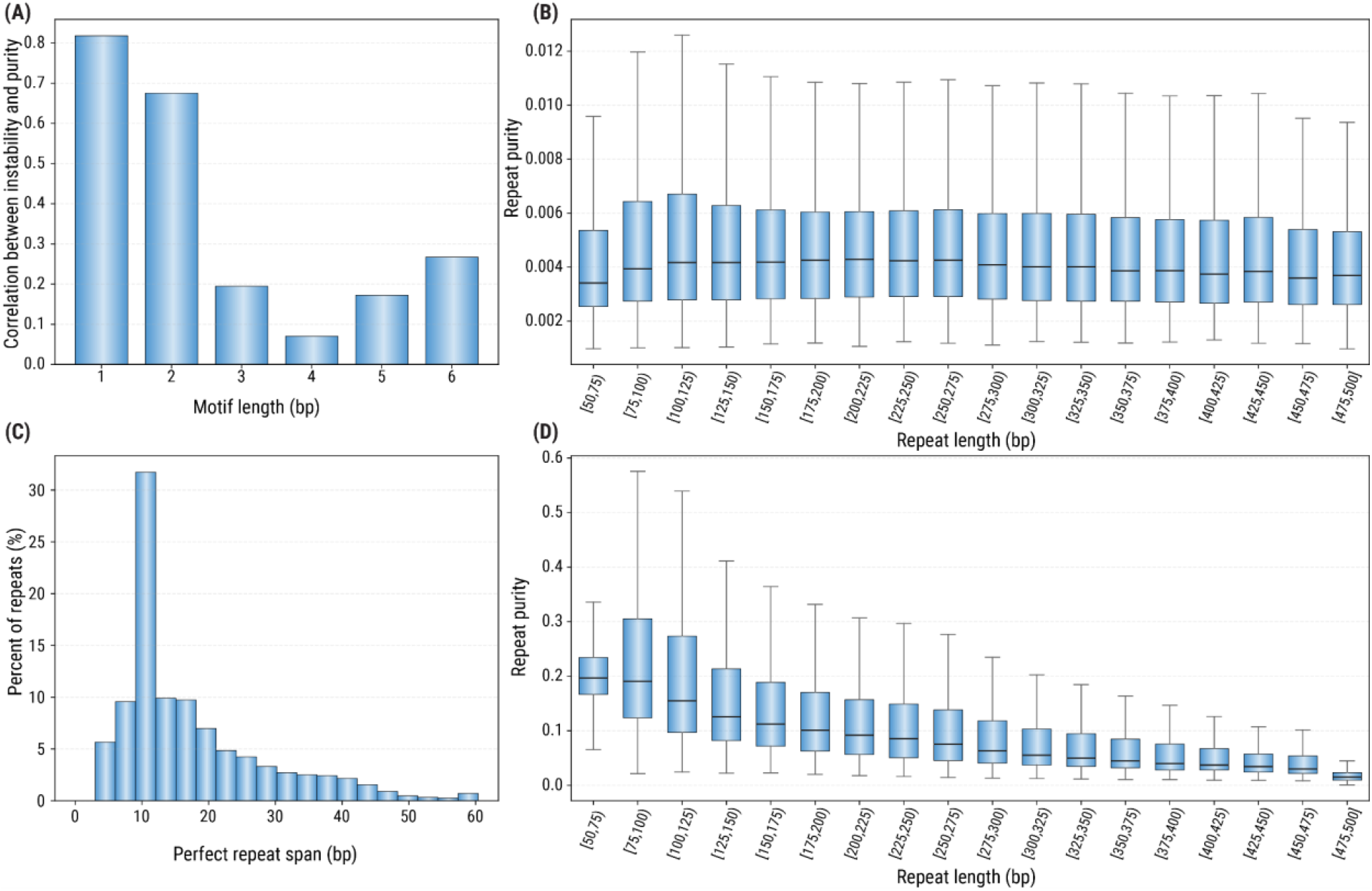
Repeat instability across HPRC. (A) Spearman correlation between repeat purity and instability rate stratified by motif length. (B) Distribution of repeat instability rates stratified by the mean repeat lengths. (C) Histogram of mean spans of perfect repeat tracts identified in repeat alleles in our catalog. (D) Mean repeat purity stratified by mean repeat length.

### Goodness of fit analysis

We assessed the goodness of fit of the DM models for 617,007 TRs with the CDF-L2 test at a p-value threshold of 0.01. When fit and evaluated on the same 256 HPRC samples, the model was rejected for 2% of loci. When the 256 samples were randomly split into two groups of 128 samples, with one group used for fitting and the other for evaluation, the model was rejected for 7% of loci. These findings suggest that the DM model describes repeat mosaicism well for most loci, while the higher rejection rate in the split-group analysis is consistent with high heterogeneity in the instability of alleles at some loci.

### Instability rates of known pathogenic repeats

We profiled instability rates of 71 known pathogenic repeat loci (**Figure 3A**) in 256 HPRC samples. The instability rate ranged from 0.002 to 0.012 across these repeats, with the fitted DM parameters ranging from tightly concentrated low-instability (e.g., *PRNP*; **Figure 3B**) to more heavy-tailed (e.g., *DMPK* and *FMR1*; **Figures 3C,D**). This locus-to-locus variability indicates that the baseline mosaicism is not uniform across repeats and suggests the need for locus-specific models. We further profiled 22 PureTarget samples with previously identified repeat expansions. We focused on 880 alleles of twenty known pathogenic repeats (*AR, ATN1, ATXN1, ATXN10, ATXN2, ATXN3, ATXN7, ATXN8OS, C9orf72, CACNA1A, CNBP, DMPK, FMR1, FXN, HTT, PABPN1, PPP2R2B, RFC1, TBP, and TCF4)*. Because mosaicism is a common feature of pathogenic repeat expansions (11,12), we assessed whether expanded alleles exhibit instability beyond the PureTarget baseline. Among 21 expansions covered by at least 10 reads, 17 were called as significantly unstable by our method (**Table S1**). This result is also consistent with our earlier observation that instability is more strongly associated with repeat purity than with repeat length, as many pathogenic expansions are known to correspond to perfect or nearly perfect repeats.

**Figure 3.**
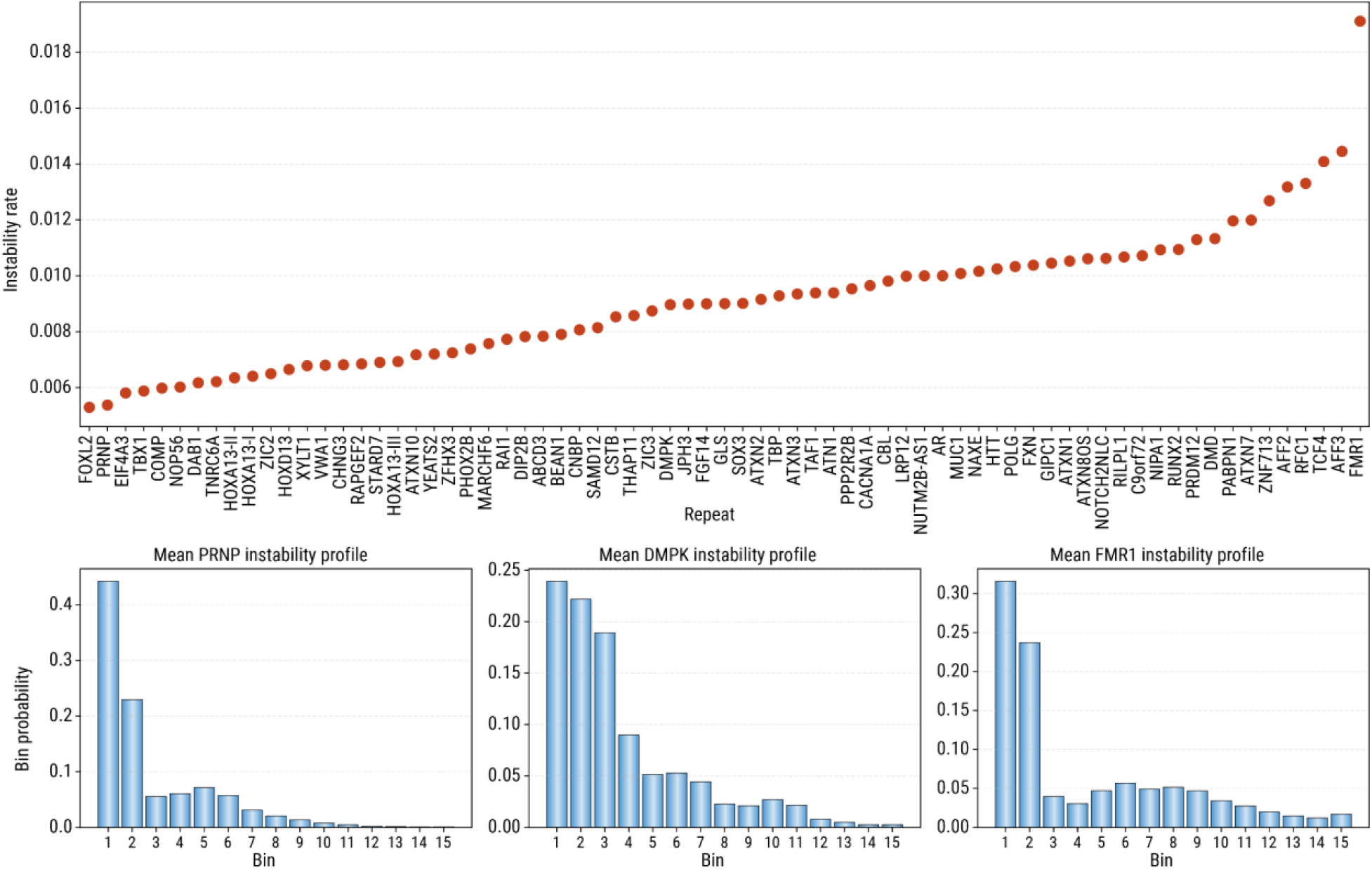
Repeat instability across pathogenic tandem repeats. (A) Instability rates of known pathogenic repeats. Fitted mean allele instability profiles for (B) PRNP, (C) DMPK, and (D) FMR1 repeats.

## Conclusions

We developed a model for quantifying TR instability from PacBio HiFi sequencing data by measuring read-to-consensus heterogeneity of repeat alleles at individual TR loci. Applied to HiFi sequencing data from 256 HPRC cell line samples, it showed that repeat instability is generally low across the genome, but varies substantially from locus to locus, supporting the need for locus-specific models rather than a single global instability measure. The model also provides a practical way to identify alleles whose instability exceeds locus-specific expectations. In particular, we found that known repeat expansions often exhibit elevated instability relative to their locus-specific baseline, suggesting that instability measures may help to prioritize candidate pathogenic repeats in genetically undiagnosed individuals.

More broadly, this work establishes a scalable, conceptually simple approach to study TR instability across both simple and structurally complex loci. Because separating true biological mosaicism from technical noise at the read level remains challenging, we do not attempt to explicitly disentangle these sources of variation. Instead, we model their combined effect to establish repeat-specific baselines, enabling robust identification of alleles with unusually high instability across samples. In this way, our method provides a practical starting point for systematic studies of repeat instability and for prioritization of disease associated repeat expansion candidates.

## Methods

### Analysis of the HPRC data

The samples from the Human Pangenome Reference Consortium were processed with the PacBio WGS Variant Pipeline v3.0.2 (13). The repeats from the adotto TR catalog v2.0 lite (see below) were genotyped in the resulting aligned BAM files with TRGT v5.0.0 (4). Subsequent repeat instability analysis was performed with TRGT-instability v1.0.0 (14).

### Read divergence rates

For each repeat locus and each sample, we extracted the alleles of the repeat and reads supporting them from the corresponding VCF and BAM files generated by TRGT. We only used alleles supported by at least 10 reads. Then we computed the edit distance between the sequence of the repeat in each read and the consensus allele. We then defined a divergence rate for each read by dividing the corresponding edit distance by the maximum length of repeat sequence in the read or the consensus. The calculation of divergence rates was implemented in the “trgt-instability divergence” command.

### Allele instability profiles

To construct allele instability profiles, we first accounted for discretization of read divergence rates. For each read, we defined a divergence-rate interval as [(k - 0.5) / L, (k + 0.5) / L], where k is the edit distance between the read and the allele consensus sequence, and L is the normalization length. We then sampled the divergence rate uniformly from this interval for each read. We partitioned the interval [0, 1] into 15 bins with the inner bin edges placed at the quantiles 0.30, 0.50, 0.65, 0.75, 0.82, 0.88, 0.92, 0.945, 0.965, 0.98, 0.988, 0.993, 0.996, and 0.998 of the empirical cumulative distribution function of these transformed divergence rates. Finally, we binned the transformed divergence rates of reads assigned to each allele into a count vector which we refer to as the instability profile of that allele.

### Defining an instability model for each repeat locus

For each repeat locus, the allele instability profiles were modeled independently with a Dirichlet-multinomial (DM) distribution *x*_*i*_ ∼ *DM*(*n*_*i*_, *α*), where *α* is the 15-dimentional parameter vector of the DM distribution, *n*_*i*_ is the sequencing read depth of allele *i*, and *x*_*i*_ is the instability profile of allele *i*. The parameter vector *α* can be expressed as *α* = *α*_0_ ⋅ *m*, where 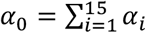 is a scalar concentration parameter and *m* = *α*/*α*_0_ is the mean instability profile. Under this parameterization, *m* describes the typical shape of the allele instability profile at a repeat locus, while *α*_0_ captures how tightly the individual allele profiles cluster around the mean profile. We estimated *m* from the aggregated binned counts, and *α*_0_ was estimated by empirical-Bayes maximum likelihood with a digamma-based Newton procedure. To account for the possibility of presence of mosaic alleles in our cohort, the model fitting was performed in two stages: We first fit the DM model using all available alleles at a repeat locus, then compute a deviance-like score for each allele, *T*_*i*_ = − *log p*_*DM*_(*x*_*i*_ |*α*_*initial*_). Alleles with values of *T*_*i*_ above the 95th percentile were considered trim candidates and at most 10% of alleles were removed. The final instability model was refitted to the retained alleles. This fitting procedure was implemented in the “trgt-instability model” command.

### A test for excess instability of an allele

To test whether a query allele was more unstable than expected, we compared the instability profile of the allele to the instability model of the repeat. The allele-level test statistic was defined as *T*_*obs*_ = − *log p*_*DM*_(*x*_*obs*_ |*α*), so larger values of *T*_*obs*_ indicate profiles that are less likely under the model. Significance was assessed by parametric bootstrap. For an observed allele with total read count of *n*_*obs*_, we simulated *n*_*sim*_ instability profiles from the instability model with the same read depths, computed the test statistic for each simulated vector, and reported the one-sided empirical p-value *p* = (1 + #(*T*_*sim*_ >= *T*_*obs*_)) / (*n*_*sim*_ + 1). We used *n*_*sim*_ = 100,000 bootstrap draws per tested allele for our analysis of PureTarget data. This test is implemented in the “trgt-instability test” command.

### Goodness of fit test

For a given repeat, we compare the observed allele instability profiles to a fitted instability (Dirichlet-multinomial) model with parameter vector *α*. We compute an empirical CDF for each observed allele instability profile by converting its counts to proportions. The expected CDF is defined from the expected multinomial probability vector *m* = *α* / ∑ *α*. The observed test statistic is the sum of squared differences between observed and expected CDF values (CDF-L2) taken over all observed allele instability profiles. To obtain the null distribution, we simulate *n*_*sim*_ datasets from the instability model while preserving the per-profile read coverages. In each simulated dataset, a multinomial probability vector is first drawn from the Dirichlet component of the instability model for each profile, then multinomial counts are sampled and converted to empirical CDFs, and the same CDF-L2 statistic is computed. The empirical upper-tail p-value is calculated as (#(simulated CDF-L2 >= observed CDF-L2) + 1) / (*n*_*sim*_ + 1). The repeat is classified as passing when p-value is above 0.01.

### Tandem Repeat Catalog

The adotto TR catalog v2.0 lite was used to define the TR loci analyzed in this study. To create this catalog, the adotto v1.2 catalog (15) was intersected with (16) to create a new set of candidate TR regions. To better centralize these regions around expected variation, read alignment from 442 long-read samples were analyzed with vclust (16). TR regions without ≥1 read having a ≥5bp insertion/deletion difference relative to the reference were filtered for appearing non-polymorphic. TR regions were additionally annotated with RepeatMasker (17) and “One code” (18) to identify and filter out tandem interspersed repeats. This combination of adotto cataloging procedure and vclust boundary centralizing is designed to capture multiple nearby and potentially interacting TRs within a single TR region. However, since known pathogenic TR loci have well defined TR boundaries, these were excluded from vclust boundary centralizing and kept at coordinates defined by STRchive (19) with a 25bp extension added. The final set of 617,007 TR regions span 86,068,019 bp (∼2.7%) of GRCh38.

## Method availability

The method described here is implemented in the TRGT-instability tool, available on GitHub (14).

## Data availability

Human Pangenome Reference Consortium data are available at the SRA under BioProject ID PRJNA730823 and the AWS Registry of Open Data: https://registry.opendata.aws/hpgp-data/. PureTarget datasets used in this study were downloaded from the PacBio datasets portal: https://www.pacb.com/connect/datasets/.

## Acknowledgements

AE and FJS receive funding from NIH (4UH3NS132105-03). We would like to acknowledge the Human Pangenome Reference Consortium (BioProject ID: PRJNA730823) and its funder, the National Human Genome Research Institute (NHGRI). GEBW is supported by an NSERC Tier 2 Canada Research Chair (CRC-2019-145/CRC-2025-00075) and an NSERC Discovery Grant Program (RGPIN-2022-04509). BID is supported by a CIHR Tier 2 Canada Research Chair (CRC-2019-00040/CRC-2023-00351) and an NSERC Discovery Grant Program (RGPIN-2022-04500).

